# A crown story: How tree architecture drives maple syrup yields

**DOI:** 10.64898/2026.06.01.728994

**Authors:** Élise Bouchard, Gauthier Lapa, Bastien Lecigne, Annie Deslauriers, Dominique Gravel, Luc Lagacé, Christian Messier

## Abstract

Maple syrup production is strongly influenced by spring weather conditions, particularly the frequency and intensity of freeze–thaw cycles. However, marked individual differences in sap and sugar yields persist among trees growing under similar stand conditions, indicating additional tree-level sources of variation. This study aimed to explain inter-individual variability in maple yields in commercial high-vacuum syrup production using structural, morphological, and growth characteristics. We used terrestrial light detection and ranging to derive variables describing crown, stem, and whole-tree size, biomass, and structure in 38 mature sugar maples. We related these variables to individual yields, including sap volume, sugar content, and syrup production. Our models explained substantial inter-individual variability: 52% in sap sugar content, 44% in sap volume, and 47% in syrup production. Laterally expanded crowns were associated with higher sap sugar content, as were lower growth rates in the first 25 mm of wood. Sap volume was highest in large, heavily branched trees with crowns extending vertically along the stem. Projected crown surface area was the strongest predictor of syrup yield, with an estimated increase of 250 mL per 5.6 m^2^. These findings highlight the importance of multidimensional crown development in maximizing individual yields in maple syrup production.

## INTRODUCTION

The spring sap of sugar maple (*Acer saccharum* Marshall) has been harvested for several centuries in the northeastern United States and southeastern Canada to produce maple syrup and various by-products, due to its particularly high sugar concentration at the end of the cold season (2 to 3.5% sugar)(Ball, 2007; Thomas, 2005). In leafless maples, sap exudation occurs during freeze–thaw events, driven by pressure changes in the wood following the formation and melting of ice, which enable the harvest of maple sap (Ceseri & Stockie, 2013; Graf et al., 2015). Thus, maple syrup yields are highly dependent on weather conditions, especially the frequency and intensity of freeze-thaw events (Bouchard et al., 2025; Tyree, 1983). However, there is also considerable variation of yields among individual maple trees growing on the same site under similar environmental conditions. Yet, the underlying causes of these inter-individual differences, especially those regarding tree architecture and vigor, remain unclear (Larochelle et al., 1998; Rademacher et al., 2023).

Maple syrup yield is determined by the harvested sap volume and its sugar content, with higher sugar content leading to similar syrup yields using less sap. However, these two components of yields have been shown to respond differently to environmental and morphological factors (Bouchard et al., 2025; Wilmot, Ellsworth, et al., 1995). Early studies investigating the relationship between diameter at breast height (DBH) and sap volume in maples have found positive relationships (Leaf & Watterston, 1964; Wilmot, Brett, et al., 1995), which was further confirmed in more recent work conducted across various sites and latitudes (Rademacher et al., 2023). However, the relationship between diameter at breast height (DBH) and sugar contents in maple sap is inconsistent, with some studies reporting positive correlations (Hauge & Lee, 1989; Wilmot, Brett, et al., 1995), while others indicating little to no relationship (Kriebel, 1990; Laing & Howard, 1990; Larochelle et al., 1998; Rademacher et al., 2023). The trunk diameter may be too coarse a metric to accurately capture inter-individual variations in soluble sugars pools and its accessibility in spring, as it does not account for the tree’s vigor nor its photosynthetic capacity.

Conversely, crown health, size, and growth rate are key long-term indicators of vigor in deciduous trees (Bossel, 1986; Moreau et al., 2020; Morin et al., 2015) and may serve as better proxies for maple sap, sugar and syrup yields. For instance, Moore et al. (2020) argued that sap volume is more closely associated with average growth rate than with DBH alone. A decline in growth due to a stress could also potentially affect sap sugar content, as stressed trees typically mobilize higher levels of carbohydrates in their tissues (Rosas et al., 2013; Tomasella et al., 2020; Wong et al., 2009), although no consistent trends have been found between growth rate and sap sweetness in sugar maple (Laing & Howard, 1990). Positive correlations between crown size and sap yield have been reported (Blum, 1971; Wilmot, Brett, et al., 1995), but Noland et al. (2006) found that sap volume were unaffected by crown dieback. The relationship between crown size and sap sugar content is also inconsistent. Some studies found no effect (Blum, 1971), while others reported a slight reduction in sugar content following crown damage (Wilmot, Brett, et al., 1995), which can persist for up to six years (Noland et al., 2006). The wide range of available metrics used to assess crown size and vigor (Schomaker et al., 2007), combined with the coarse resolution of ground-based measurements that cannot capture fine-scale architecture, such as proportion and size of branches, may explain the observed inconsistencies.

Indeed, low order branches (i.e. smaller diameter) likely influence sap exudation, as their low volume-to-surface ratio causes them to thaw and freeze first under shifts in solar radiation and temperatures. In line with this hypothesis, sap flow rates in leafless maples have been shown to correlate more closely with temperature changes in the branches of the canopy than with those in the bark or in the xylem (Marvin & Erickson, 1956; Plamondon & Bernier, 1980). Greater wood surface area in contact with air may therefore increase yields by enhancing freeze–thaw dynamics. Yet, a full assessment of maple architecture in relation to sap yields, across branches of all orders, has never been conducted, likely due to the labor-intensive nature of measuring these canopy components. However, light detection and ranging (LIDAR) now enable detailed assessments of tree architecture, as well as more accurate estimates of traditional tree size metrics (Srinivasan et al., 2015).

In addition to tree architecture, harvesting systems significantly affect the sap volume collected from trees. Modern commercial maple syrup production utilizes vacuum pumps and tubing systems, which can yield up to five times more maple sap per tap relative to traditional gravity systems using buckets where sap is passively harvested through gravity (Wilmot et al., 2007). Nevertheless, except for Moore et al. (2020), all studies cited above examining the effects of tree morphology and growth on maple syrup yield relied on the traditional bucket collection method. Consequently, it remains unclear whether these findings apply to commercial operations.

The objective of this work was to provide a comprehensive assessment of how tree architecture and growth rate impact maple syrup yields and its two components, sap volume and its sugar content (Brix), under commercial high-vacuum practices. Traditional tree size and growth metrics (DBH, tree-ring width, height, and crown diameter) were measured alongside more complex structural and biomass variables derived from terrestrial LIDAR, including crown volume, projected surface area, and asymmetry, as well as wood surface area and volume across all axes (main stem and branches). We aimed to identify the main drivers of inter-individual variability in maple syrup production acting at the stand level, thereby providing producers with practical insights into implementable management practices. We hypothesized that crown size metrics would be stronger predictors of maple syrup yield and its components than traditional stem size variables (e.g., DBH and tree height), due to their closer link to tree vigour and key physiological processes, including spring pressure generation in wood and carbohydrate dynamics.

## MATERIAL AND METHODS

### Study site

This experiment was conducted at the experimental sugar bush of the Acer Research Center in Saint- Norbert-d’Arthabaska, Québec, Canada (1.4607646° N, 71.8871713° W). The site receives an average of 1,940 hours of sunlight annually, 1,091 mm of precipitation (810 mm rain, 280 mm snow), and has a mean temperature of 4.5 °C. The stand is primarily composed of sugar maple (*Acer saccharum Marsh*.), with a low proportion of American basswood (*Tilia americana L*.), American beech (*Fagus grandifolia Ehrh*.), ironwood (*Ostrya virginiana (Mill*.*) K. Koch*), and occasional yellow birch (*Betula alleghaniensis Britton*). Eastern hemlock (*Tsuga canadensis (L*.*) Carrière*) and eastern white cedar (*Thuja occidentalis L*.) occur locally in topographic depressions. The average age of the stand is 110 years, with a potential of 1800 taps for the 17.2 ha forest.

### Tree selection

Extensive branch damages caused by a major ice storm in January 1998 led to marked variation in crown size and architecture across the site. In autumn 2019, trees were visually preselected to capture this range of crown structures, as well as variation in previous-year tap closure rates, used as a proxy for growth rate. A total of 38 co-dominant or dominant sugar maples were selected, with diameters at breast height (DBH) ranging from 40 to 62 cm, averaging 49 cm, and total heights between 23 and 28.5 m, averaging 26 m.

### Sap collection

Trees were tapped at the beginning of March 2020, with each tree receiving only one tap. Taps were drilled to a depth of 4.45 cm (1¾ in) and a diameter of 7.94 mm (5/16 in), while maintaining a horizontal spacing of 15–25 cm (6–10 in) from the previous year’s taps. For each tree, sap was collected in a stainless-steel barrel with a capacity of 205 L (45 US gallons), connected to a vacuum pump. The barrel was sealed to maintain internal vacuum pressure. Vacuum levels throughout the sugaring season ranged from 25 to 28 inHg, corresponding to approximately 85–95 kPa. The barrels were carefully sanitized before installation in the forest to prevent microbial contamination and sugar degradation during the season, following the protocol described in Appendix S1. At the end of the sugaring season (April 15), the barrels were transported using an excavator, after which the sugar content (Brix) and mass of the sap in each barrel were measured using a refractometer and an electronic balance, respectively. The sap volume (*V*_*sap*_, L) was calculated by dividing the sap mass (*M*_*sap*_, Kg) by the sap density (*D*_*sap*_, kg · m^-3^), with sap density determined based on the measured sugar content according to equation 1 and 2 from Grenier (2007), where *Brix*_*sap*_ is the measured Brix of the sugar sap.

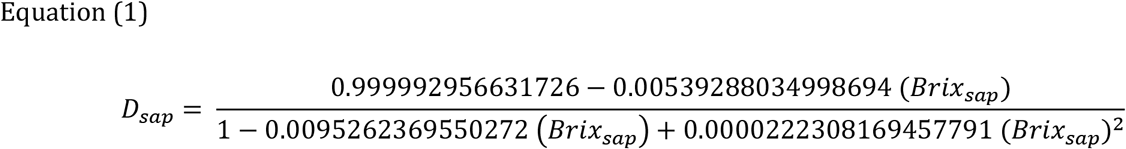

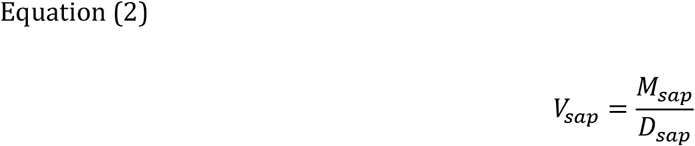

Both *D*_*sap*_ and *V*_*sap*_ were used to compute potential syrup yield (*V*_*syrup*_, L), according to equation 3, where the Brix value of the syrup (Brix_syrup_) was set at 66 ° Bx with a density of 1.3248.

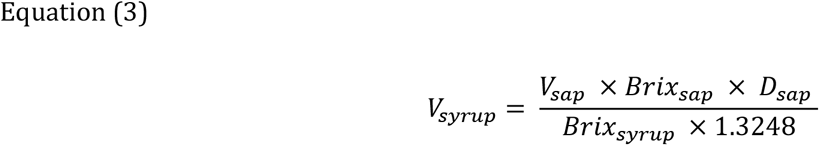

### Radial growth

During the summer of 2021, a core was sampled from each tree using a 5.15 mm increment borer (Haglöf, Mora, Sweden), taken 50 cm below the tap. The cores were glued onto a wooden molding, oven-dried at 60 °C until reaching constant weight and finely sanded. Ring widths were measured using a Velmex system (Velmex Inc., Bloomfield, NJ, USA) with an accuracy of 0.001 mm. Average growth rates were estimated at depths of 25 and 75 mm, excluding the bark. Four cores were defective and could not be used, leaving a total of 34 trees with usable growth data.

### Tree laser scanning

Selected trees were scanned with a Terrestrial Laser Scanner (TLS) Focus 3D X130 (FARO Technologies, Inc., Lake Mary, Florida) in November 2020, following leaf drop and prior to the first snowfall. Three to four scans were captured around each tree to ensure full coverage, with georeferenced targets placed around the tree to accurately register and align the multiple scans. Scans were then registered and combined using Faro Scene (version 5.3.3.38662). Point clouds were manually cleaned in CloudCompare (version 2.11.1). Each tree was processed individually, and all points not belonging to the target tree were removed.

### Tree metrics

Individual tree point clouds were denoised using the *VoxR* package (Lecigne, 2020), whereby points with a mean distance to their 8 nearest neighbours exceeding the cloud-wide mean by more than 0.5 standard deviations were removed as noise. Quantitative structure models were constructed with the aRchi package (Martin-Ducup & Lecigne, 2025), and branch orders were assigned, with the stem defined as order 1. Three structural categories were defined: total, representing the entire tree; main axis, including branch orders ≤ 2; and branches, including orders ≥ 3. These categories were consistently used to partition both wood surface area (m^2^) and wood volume (m^3^). Functions from *aRchi* were used to estimate wood surface area for each category, as well as the fork rate (*n* forks · m^-1^) at the whole tree level. The *VoxR* package was subsequently used to voxelize the point cloud and estimate wood volume for the same structural categories, and to compute crown diameter using the distant-point method. Finally, the *ITSMe* package (Terryn et al., 2023) was used to estimate crown volume (m^3^) and projected surface area (m^2^) based on concave hulls, and to derive tree height (m) and diameter at breast height (DBH, cm).

Custom functions were developed to derive additional architectural metrics. Tree inclination was quantified as the horizontal displacement between the centroids of stem slices at breast height (DBH) and at the crown base, normalized by the distance between these two centroids along the stem axis. Stem slices were extracted using the *stem_slice()* function from the *ITSMe* package (Terryn et al., 2023). Centroids for each slice were estimated using a circle-fitting approach, while crown base height was defined as the highest Z-coordinate of trunk points classified with *ITSMe*, minus a 60 cm buffer to avoid slicing within forks. For trees with forks extending below this threshold, the buffer was manually adjusted. Crown asymmetry was computed following Franco (1986), identified as one of the most robust crown asymmetry indices in trees (Kong et al., 2021). The method is based on a crown displacement vector within a circular reference framework, which we derived from the summed XY coordinates of radii measured in 36 azimuthal directions around the crown centroid (5° angular tolerance). In each direction, radius length was defined as the 95th percentile of point distances. The crown-base centroid was computed as the geometric center of a convex hull constructed around the trunk points at the crown base identified by *ITSMe*, after applying a 60 cm buffer. Crown length (m) was calculated as the difference between total tree height (m) and the height of the lowest crown points, classified using *ITSMe*. All variable names, descriptions and units are listed in Table 1.

**Table 1.**
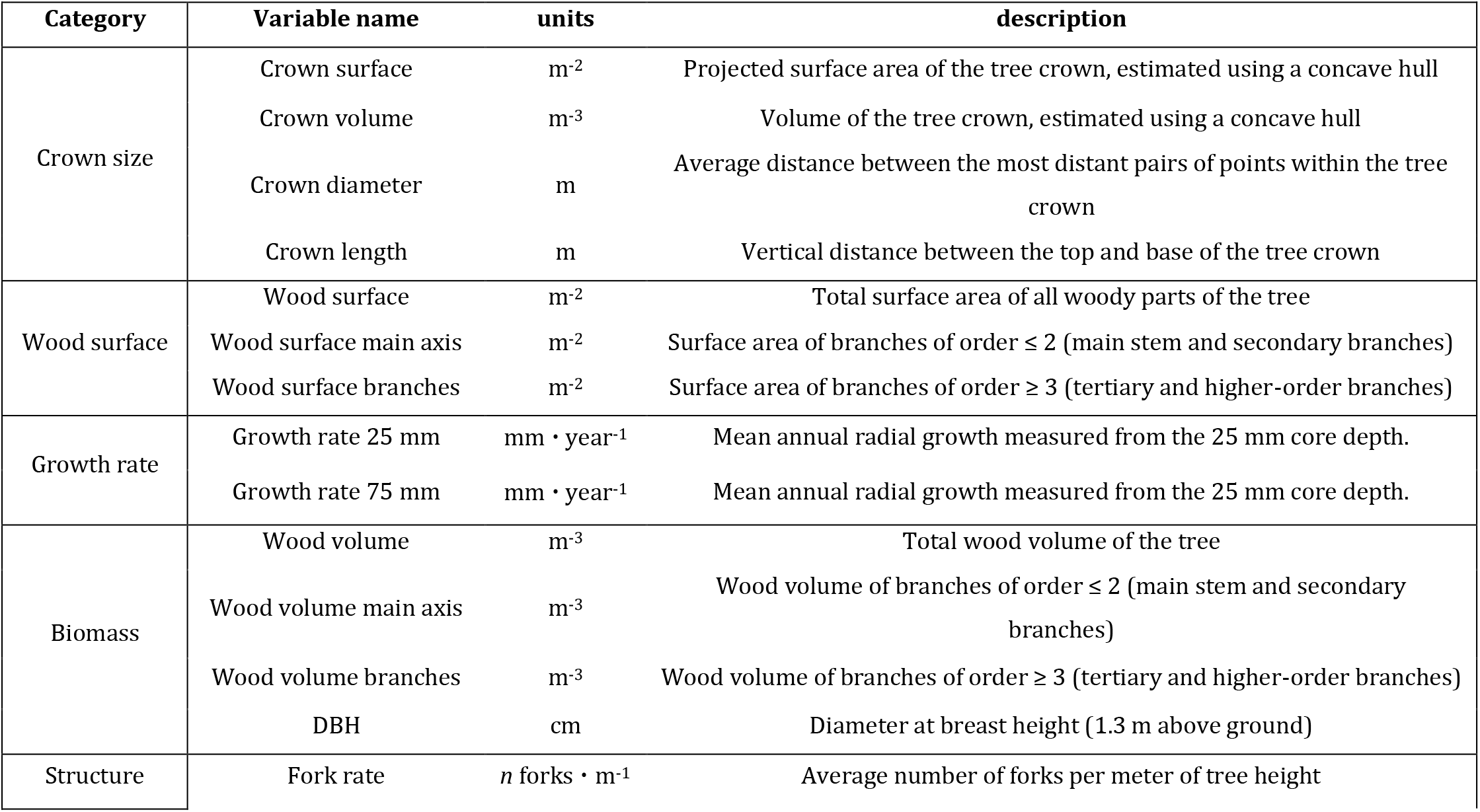
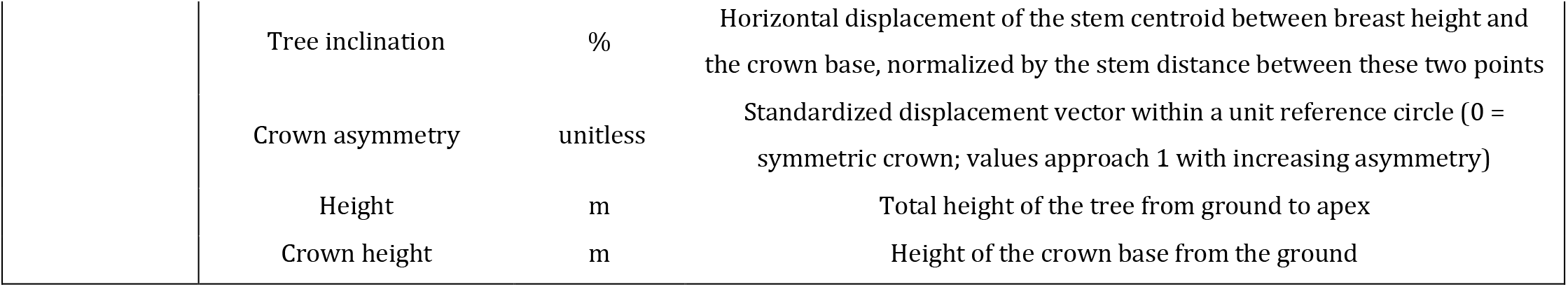
Categories, names, descriptions, and units of all tree size, architecture and growth variables, derived from terrestrial laser scanning or tree-core measurements.

### Statistical analyses

Pairwise relationships (bivariate linear regressions and Pearson correlations) were first examined to provide a rapid assessment of associations among variables and to guide data screening and cleaning. Principal component analysis (PCA) was then conducted to explore the main axes of variation among predictors. Multiple linear regressions were fitted to model relationships between yield response variables (sugar content, sap volume, and maple syrup volume) and morphological and growth variables. Multiple candidate models were developed for each yield component as response variable. Model construction was informed by prior knowledge from the literature on factors influencing yield components, allowing us to test biologically grounded hypotheses. Results from pairwise and PCA analyses guided model construction, ensuring that only variables contributing new information were included, while redundant variables were avoided. This approach also helped limit collinearity, which was checked using the variance inflation factor (VIF), from the package *car* (Fox et al., 2012), with a conservative threshold of 3. Model selection was performed using the Akaike information criterion corrected for small sample size (AICc) using the package *AICcmodavg* (Mazerolle & Mazerolle, 2017). Models with ΔAICc ≤ 3 were considered equivalent. Model assumptions were assessed using the *DHARMa* package (Hartig & Hartig, 2017), which simulates standardized residuals to evaluate heteroscedasticity, residual normality and outliers. Cook’s distance was additionally used to assess the influence of individual observations, with values greater than 1 indicating highly influential points.

### Rationale for the models

The rationale underlying model construction is summarized below, while a full description of all candidate models and the model-selection procedure presented in Appendix S2.

#### Sap sugar content

Sugar maples tend to store more carbohydrates in crown branches than in the trunk at springtime (Wong et al., 2003), and crown structure and health reflect long-term tree vigour and photosynthetic capacity in deciduous trees (Moreau et al., 2020; Morin et al., 2015). In addition, maple sap is largely sourced from nearby tissues (Bouchard et al., 2025), and stressed trees often exhibit increased carbohydrate mobilization (Rosas et al., 2013; Tomasella et al., 2020; Wong et al., 2009).

Based on these evidences, we expected sap sugar content to be primarily influenced by crown size and structure, with effects potentially modulated by proximity to the crown and by physiological stress, leading to the following hypotheses: (i) a **Crown hypothesis**, testing different metrics of crown and branch size; (ii) a **Distance hypothesis**, evaluating the effect of proximity to the crown after accounting for overall crown size; (iii) a **Stress hypothesis**, where reduced recent growth serves as an indicator of physiological stress; and (iv) a **Shape hypothesis**, incorporating architectural traits such as crown asymmetry and fork rate to assess their role in soluble sugar mobilisation.

#### Sap volume

Larger trees (as indicated by DBH) tend to produce more sap (Leaf & Watterston, 1964; Rademacher et al., 2023; Wilmot, Brett, et al., 1995). Sap exudation is also driven by freeze–thaw cycles within the conductive system (Tyree, 1983), which may be enhanced by greater woody surface area exposed to ambient air. In addition, larger volumes of conductive wood (sapwood) are expected to store and mobilize more sap, consistent with the pipe model theory, which links sapwood area to crown-supported branch biomass (Lehnebach et al., 2018).

Based on these facts and mechanisms, sap volume was expected to increase with greater wood volume, woody surface area, and greater branching along the trunk. We therefore formulated five hypotheses: (i) a **Wood volume hypothesis**, testing the effects of DBH and total/main-stem wood volume on sap availability; a **Pressure hypothesis**, testing woody surface area, which may affect sap flow via hydraulic pressure shifts from enhanced freeze–thaw cycles; (iii) a **Combined hypothesis** including both wood volume and surface area, assuming additive effects on sap yields; (iv) a **Growth hypothesis**, where growth rate may indirectly influence sapwood area, thereby modulating sap yields; and (v) a **Branch hypothesis**, testing whether greater branch volume along the trunk is associated with higher sap volume, possibly to support a greater crown hydraulic demand.

#### Maple syrup

Because maple syrup yield depends on both sap sugar content and sap volume, combining the variables that best predict each component should allow accurate prediction of syrup yield. Accordingly, the syrup yield models were based on four hypotheses: (i) a **Single component** hypothesis including the best predictors of either sugar content or sap volume; (ii) a **Combined** hypothesis including predictors of both components; a **Reduced** hypothesis excluding potentially redundant variables; and (iv) a **DBH-only control**, included because DBH is widely measured in forestry and allows comparison with simpler, commonly used models.

## RESULTS

### Yields

Total sap volume collected per tree over the season ranged from 51.0 to 148.8 L (mean of 98.4 L), with sugar content varying between 2.2 and 4.2 °Bx (mean of 3.0 °Bx). Potential maple syrup yield ranged from 2.1 to 5.7 L (mean of 3.5 L). Sap volume and sugar content were not correlated (adj. R^2^ = −0.02), confirming that these two components are likely not influenced by the same environmental and morphological constraints. Maple syrup yield was strongly influenced by both components (sap sugar content adj. R^2^ = 0.37; sap volume adj. R^2^ = 0.57), indicating that accurate estimation of both is required for efficient prediction of maple syrup yields under high-vacuum commercial systems.

### Structure of Morphological Variation

DBH was modestly correlated with crown dimensions and biomass but was a poor correlate of growth rates and crown length (Fig. S3.1). Its strongest correlation was with total wood volume (r = 0.56). Trees with larger trunks also tended to have bigger crowns; however, the low r pearson for crown dimension variables (r = 0.09 to 0.47) indicated that DBH was not a strong proxy for estimating crown size in these mature sugar maples. Its intermediate position along the two main PCA axes confirmed this ambiguous role (Fig. 1). Growth rate, both in recent years (25 mm increment) and over past decades (75 mm increment), was associated with crown size variables, especially with those acting on a vertical axis such as crown length and crown volume (r = 0.43 to 0.62, Fig. S3.1).

**Fig. 1.**
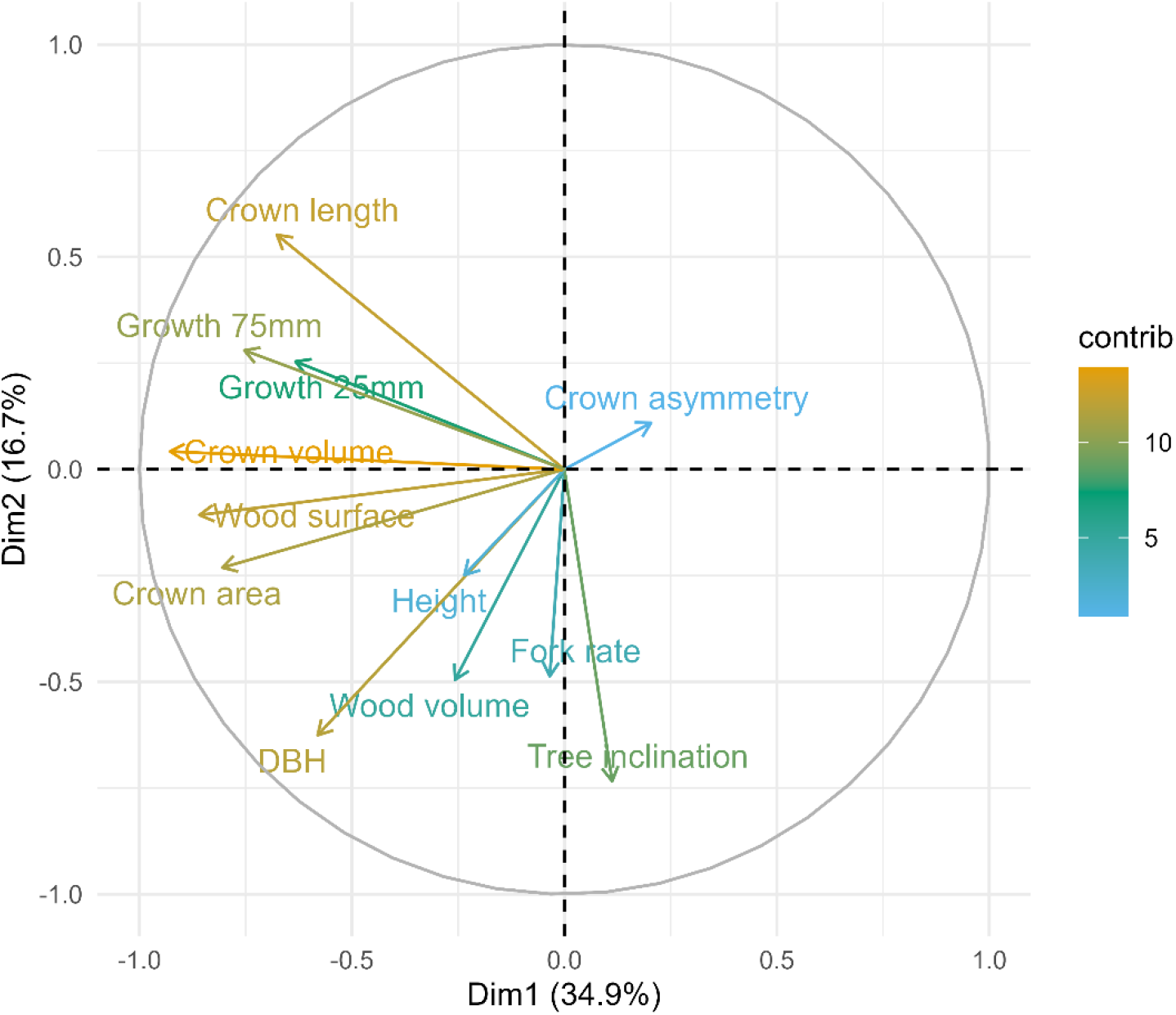
Principal Component Analysis (PCA) of tree architecture and growth rates. The biplot displays the two main axes of variation, illustrating covariation among stem size (DBH, wood volume), crown dimensions (crown volume, crown length, crown surface area, wood surface), structural traits (crown asymmetry, tree inclination, fork rate, height), and growth rates. Arrows indicate the direction and relative contribution of each variable along the principal components. Percentages in parentheses next to each axis represent the proportion of total variation explained by that component.

All crown dimension variables, including crown surface area, crown volume, branch wood volume, woody surface area, and crown length, were positively associated with one another (r = 0.38 to 0.86). The only exception was crown asymmetry, which showed weak negative correlations with branch volume, woody surface area, and both crown volume and surface area (r = -0.19 to -0.42). Trees with asymmetric crowns tended to exhibit smaller crowns and reduced branch volume. This explains why the first PCA axis, which accounted for 34.9% of the variation, was largely dominated by crown-related variables, contrasting trees with extensive crowns, high growth rate, and substantial woody surface area on the left, with trees having poorly developed crowns on the right and low growth rate (Fig. 1).

Height was an independent variable in the architecture of the trees, with very few and weak correlations with other variables, excepting for DBH (r = 0.47). Forking rate was poorly associated with any other variables, indicating that forks might occur in a vast array of stem and crown sizes. Tree inclination was also weakly correlated with most variables, although more inclined trees tended to have crowns positioned higher along the stem, resulting in shorter crown length (r = -0.50).

Understanding the pairwise relationships and axes of variation among structural attributes of maple trees guided the construction of yield prediction models by selecting complementary variables and avoiding redundant ones that could introduce collinearity. The data will be publicly available on Figshare upon publication.

### Sugar content

The best model predicting sugar content was Model 7 (Stress hypothesis), which clearly outperformed the second-best model (Model 4, Shape hypothesis; ΔAICc = 11.04; Table S2.2). This model explained 52% of the variance (adjusted R^2^). All three predictors (crown area, growth at 25 mm depth, and main stem wood volume) were statistically significant (p < 0.05). An increase of 1 m^2^ in crown area was associated with a 0.03 °Bx increase in extracted sap sugar content (SE = 0.005; Fig. 2). Sugar content also increased by 0.31 °Bx (SE = 0.11) per 1 m^3^ increase in main stem wood volume, whereas higher recent growth (averaged over 25 mm depth) was associated with a decrease of 0.60 °Bx per mm·yr^−1^ (SE = 0.14). Therefore, trees with crowns covering a large area, high wood biomass in the main stem, and low recent growth rates tended to have sweeter sap.

**Fig. 2.**
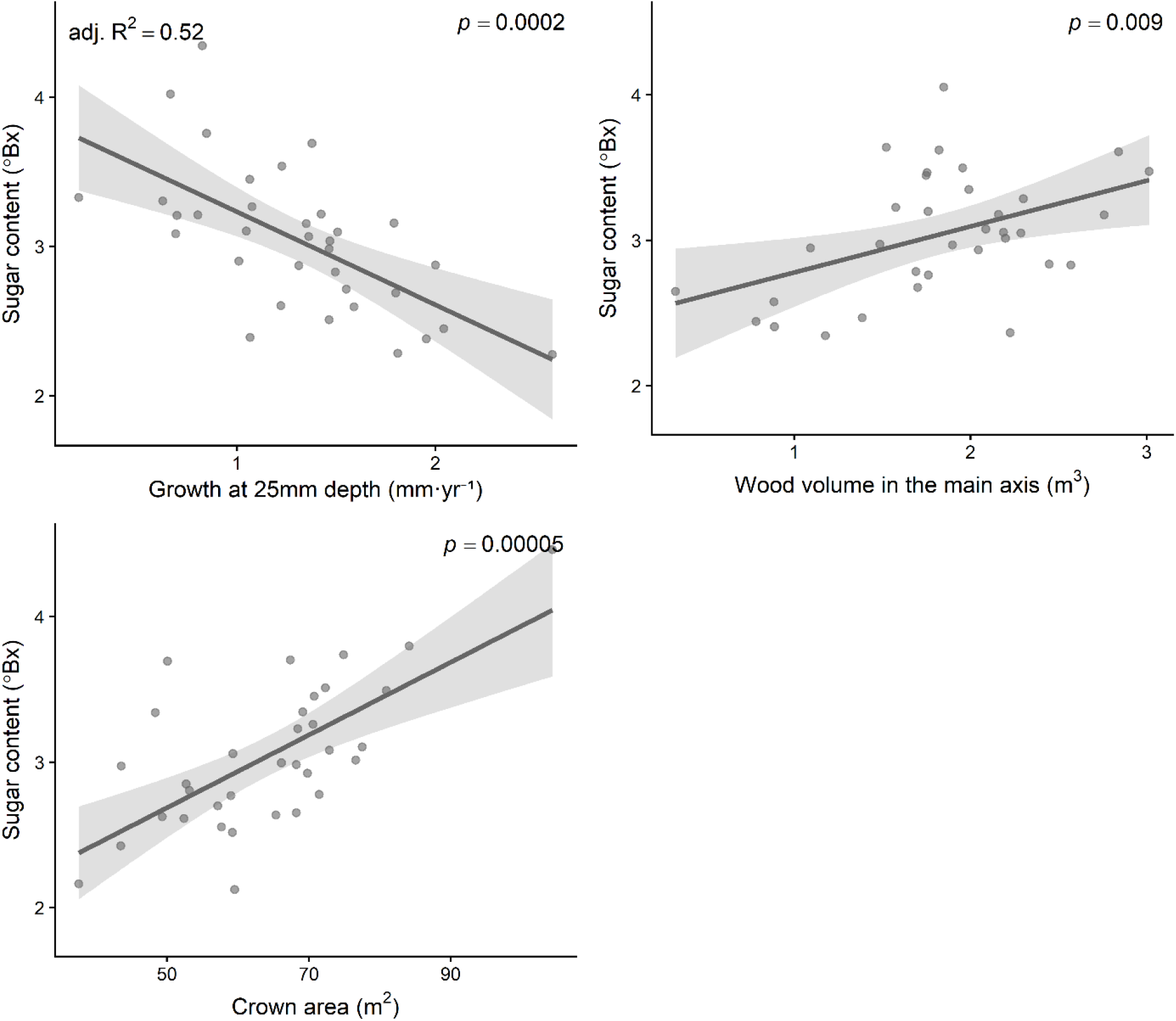
Effect plots from the best-fitting model predicting sap sugar content (°Bx) as a function of growth at 25 mm depth (mm·yr^−1^), main axis wood volume (m^3^), and crown area (m^2^). Points represent partial residuals, and shaded bands indicate 95% confidence intervals. Model fit statistics (adjusted R^2^ and p- values) are displayed on the plots.

### Sap volume

Models 9 and 8 for predicting sap volume, both related to the branch hypothesis, were similarly supported (ΔAICc = 1.3; Table S2.4). Model 8 was more parsimonious, including only two predictors (crown length and DBH) and yielding an adjusted R^2^ of 0.37, whereas Model 9 included two additional predictors and achieved a higher explanatory power (adjusted R^2^ = 0.44). Although Model 8 would be preferred for purely predictive purposes due to its greater parsimony, Model 9 was selected based on its stronger biological interpretability, as it provided a more detailed description of how the allocation of woody biomass (branch volume and main stem volume) influences sap production. The selected model (model 9, branch hypothesis) explained 44% of the variance (adjusted R^2^). All effects were statistically significant (p < 0.05), except for wood volume in the main axis, which was marginally significant (p = 0.05). Sap volume increased with crown length (3.19 ± 1.18 L m^−1^), branch wood volume (104.83 ± 45.56 L m^−3^), and DBH (2.15 ± 0.68 L cm^−1^), but decreased with wood volume in the main axis (−13.75 ± 6.76 L m^−3^, Fig. 3). Together, these results indicate that larger trees with vertically extended crowns and a greater allocation of wood to branches relative to the main stem tend to produce more extractable sap volume during spring.

**Fig. 3.**
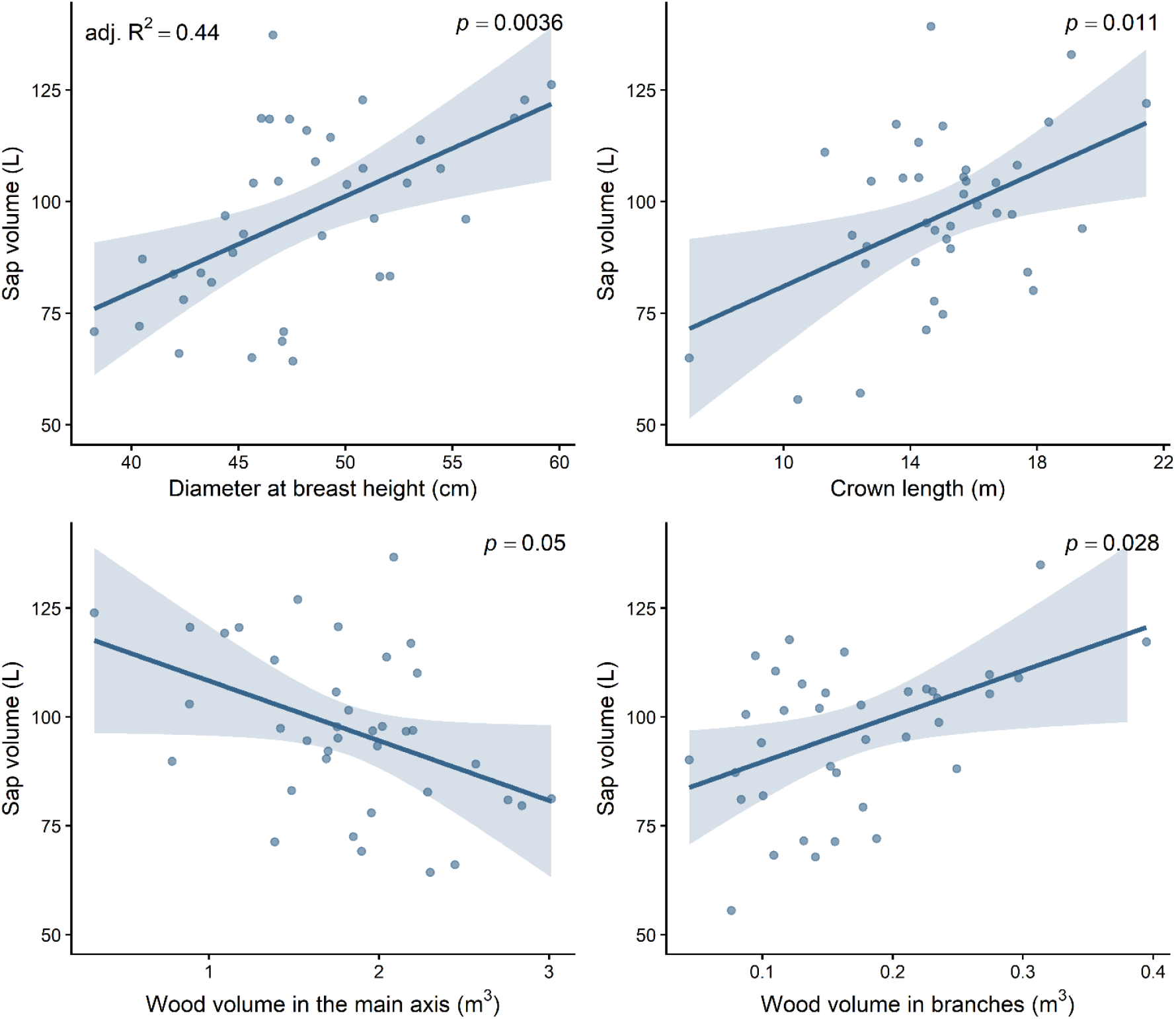
Effect plots from the selected model predicting sap volume(L), including four predictors: diameter at breast height (cm), wood volume in the main axis (m^3^), wood volume in branches (m^3^), and crown length (m). Points represent partial residuals, and shaded bands indicate 95% confidence intervals. Model fit statistics (adjusted R^2^ and p-values) are displayed on the plots.

### Maple syrup

Models 5, 4, and 3, corresponding to the Combined and Reduced hypotheses explaining maple syrup yield, were well supported (ΔAICc ≤ 1.03; Table S2.6). The most parsimonious model (model 5), which explained 45% of the variance, included crown area as the sole explanatory variable but exhibited heteroscedastic residuals, particularly overpredicting the lowest syrup yields. Consequently, we selected model 6 as the best candidate model, which added DBH as a second explanatory variable. Although this additional term was not statistically significant, it reduced heteroscedasticity in the residuals.

The selected model (model 4, Reduced hypothesis) explained 47% of the variance (adjusted R^2^). Crown area had a significant positive effect on maple syrup yield (p < 0.05), with yields increasing by approximately 0.05 L per 1 m^2^ of crown surface area (SE = 0.010 L m^−2^). The effect of DBH was not significant, indicating limited additional explanatory value after considering crown area. Its estimated effect corresponded to an increase of 0.04 L per cm of DBH (SE = 0.03 L cm^−1^, Fig. 4).

**Fig. 4.**
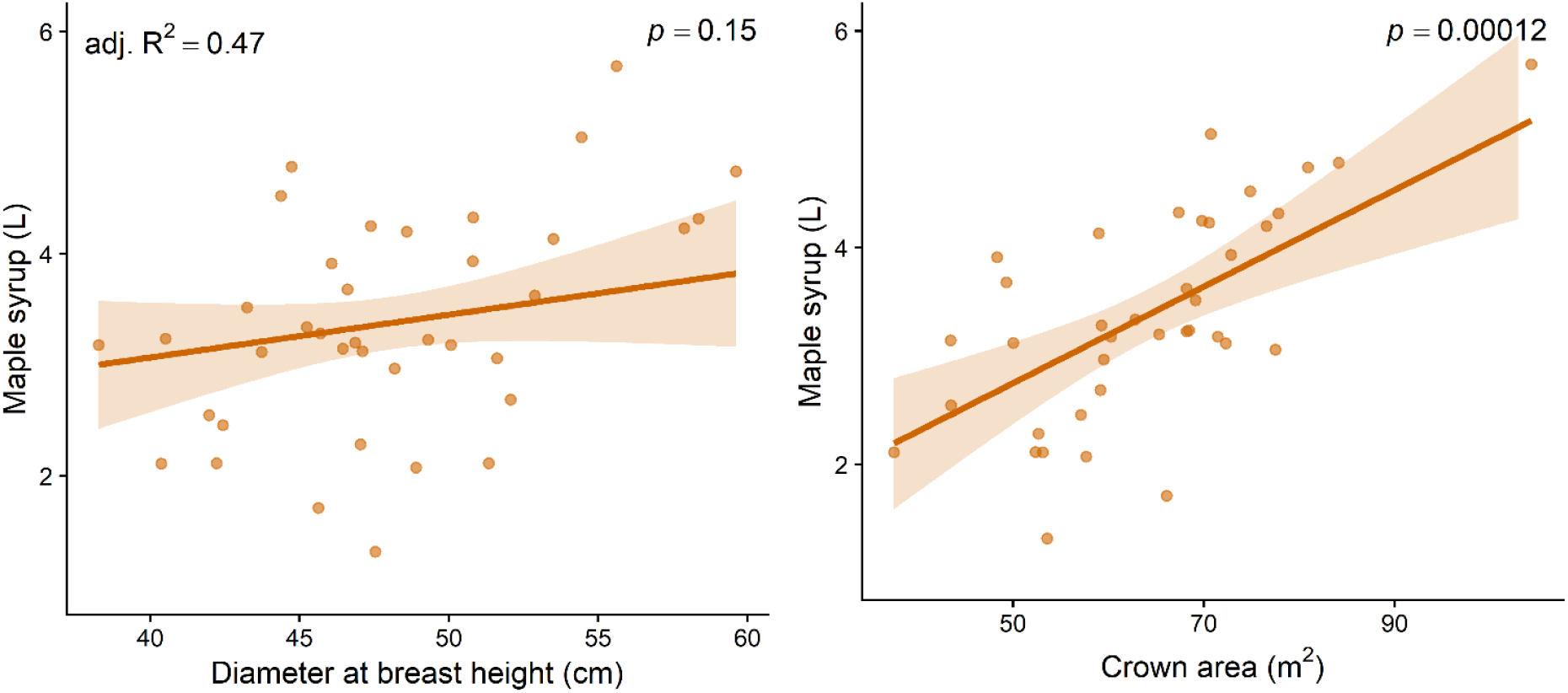
Effect plots from the selected maple syrup yield (L) model with two predictors: diameter at breast height (cm) and crown area (m^2^). Points represent partial residuals, and shaded bands indicate 95% confidence intervals. Model fit statistics (adjusted R^2^ and p-values) are shown on the plots.

Variance inflation factors were below 1.90 for all predictors across all selected models, indicating negligible collinearity among predictors.

## DISCUSSION

Consistent with our hypotheses, variation in crown size and structure among neighboring trees strongly contributed to differences in maple syrup yield and its components over a production season. Sugar maple is among the most shade-tolerant hardwoods in northeastern North America (Burns et al., 1990), a trait often associated with strong crown plasticity to enable adaptation to low and heterogeneous light conditions (Valladares & Niinemets, 2008). Accordingly, sugar maple has been shown to develop long lateral branches at different heights to optimize light capture in response to canopy gaps or the proximity of competitors (Brisson, 2001; Martin-Ducup et al., 2016), making sugarbushes naturally prone to a wide diversity of crown sizes and shapes.

We acknowledge that these relationships may differ in younger trees (<40 cm DBH), as crown morphological differences in sugar maple emerge late in ontogeny, being pronounced in 55-year-old stands but absent in 35-year-old stands, in both mixtures and monocultures (Martin-Ducup et al., 2016, 2018). Indeed, crown-shaping processes, such as gap dynamics and branch damage, operate and accumulate over long-time scales. Dominant sugar maples have typically experienced multiple gap-forming events before reaching the canopy (Canham, 1985), and twig mortality due to crown abrasion is greater in larger trees (Hossain & Caspersen, 2012), both processes potentially generating variation in crown size and architecture that may influence sap and syrup yields.

Traditional forestry metrics such as DBH and tree height appeared to be less closely linked to maple syrup yield and its individual components (sap sugar content and sap volume). Tree height did not influence any yield component, whereas DBH was associated with greater sap volume but not with sap sugar concentration, consistent with results from previous studies (Rademacher et al., 2023). However, once crown size was considered, DBH contributed little additional information for explaining syrup yield. These contrasting relationships may help explain inconsistencies reported in the literature regarding the influence of tree size on yields (Blum, 1973). Moreover, growth rate averaged over 25 or 75 mm depth did not seem to perform better than DBH on predicting sap volume, contrary to previous findings (Moore et al., 2020). Overall, crown size metrics emerged as the most promising proxies for capturing inter-tree variation in maple syrup yields and its individual components (sap volume and sugar content) in mature sugarbushes under high-vacuum production systems.

### SAP SUGAR CONTENT

Explaining half of the variation among trees in sap sugar content is notable, given the still poorly understood drivers of sap sugar content at springtime (Hauge & Lee, 1989; Kriebel, 1990; Laing & Howard, 1990; Larochelle et al., 1998). Larger crowns would be expected to support higher sap sugar content, as they represent large pools of soluble sugars and major sinks in spring (Martínez-Vilalta et al., 2016; Wong et al., 2003). Our results support this expectation, although it remains unclear why the effect is not observed for total crown volume but is instead driven by horizontal crown attributes (area and asymmetry), which largely reflect the presence and distribution of long lateral branches around the stem. In fruit orchards, horizontal branches enhance fruit quality, yield, and photosynthetic performance relative to upward- or downward-oriented branches (Yang et al., 2019), potentially creating stronger spring sinks for soluble sugars. The role of such a mechanism in maple syrup production has yet to be determined, and a more detailed analysis of tree architecture, particularly branch angle, could help elucidate this relationship. Greater wood volume in the stem and secondary branches was also associated with sweeter sap after accounting for crown size and recent growth. The main stem stores a large proportion of non-structural carbohydrates in temperate trees (∼40%; Barbaroux and Bréda 2002; Richardson et al. 2015), and soluble sugars from this pool are readily accessible during maple syrup production because harvested sap is mainly drawn from tissues near the tapping zone (Bouchard et al., 2025). Thus, larger stem volume seem to increase spring sugar availability in sap.

Recent growth in wood rings, averaged over 25 mm depth, was the strongest predictor of sap sugar content, with sweeter sap associated with lower recent growth rates, potentially reflecting a recent stress event in the tree’s life. Indeed, trees commonly mobilize non-structural carbohydrates in response to stresses such as drought or cold (Rosas et al., 2013; Tomasella et al., 2020; Wong et al., 2009). Consequently, we initially formulated this hypothesis as a stress hypotehsis. However, recent low growth rate may also reflect ontogenetic changes in mature trees, as the study was conducted in an old stand (∼110 years old). In late developmental stages, shoot elongation and radial growth naturally decline, with carbon allocation shifting from spatial exploration toward light capture (Taugourdeau et al., 2019) and reproduction (Millet, 2012). This natural shift in carbon allocation could come with increased sugar levels in spring sap. Therefore, retaining large, old trees in maple syrup productions could be beneficial, not only because their size provides sufficient wood volume for sustainable tapping (Messier et al., 2022), and supports habitat availability for biodiversity (Cadieux et al., 2023; Martin et al., 2022), but also because sap sugar content may naturally increase as the tree matures. There was no clear evidence of recent stress in the growth-ring series; however, it was not possible to disentangle stress from ontogeny as drivers of growth decline, as heartwood decay in many trees resulted in incomplete series, limiting the reconstruction of long-term ontogenetic trends and stress history. A larger sample of growth series spanninng the full-lifespan of mature sugar maples would help disentangle these two potential causes of growth decline, which in turn strongly impact sap sugar content.

The effect of reduced growth on increased sugar content was evident at shallow depth (25 mm) but absent at 75 mm, suggesting that only recent growth rings were influential. Several lines of evidence suggest that outer growth rings play a key role in determining spring sap sugar concentration. For example, Perkins et al. (2021) showed slightly higher sugar concentrations in shallow taps (25 mm) compared to deeper taps (38–64 mm), suggesting higher sugar availability near the bark. Similarly, Essiamah and Eschrich (2014) found that spring sap-producing species (birch, maple, alder) have lower starch levels in bark parenchyma than in wood at springtime, unlike non-exuding species (beech, oak, ash), indicating that soluble sugars derived from starch hydrolysis may be mobilized near the bark. In temperate hardwoods, carbohydrate in outer growth rings (0–1 cm) are the most dynamic, showing strong seasonal variation and dominating metabolic fluxes whereas deeper reserves are rarely mobilized (Furze et al., 2018, 2021; Hoch et al., 2003; Zhang et al., 2014), a pattern also observed in spring maple sap, where soluble sugars mainly originate from recent carbon pools integrating three to five years on average (Muhr et al., 2016).

Together, these findings suggest that early-spring sap sugar dynamics are closely linked to activity in the outer growth rings near the bark. In contrast, sap volume does not follow this pattern, as it can increase with tapping depths up to at least 50 mm in vacuum systems (Perkins et al., 2021).

### SAP VOLUME

A greater number of branches along the trunk (longer crown length) for a given DBH was associated with higher sap volume, consistent with allometric rules governing tree architecture, including the pipe model and Leonardo da Vinci’s rule (Lehnebach et al., 2018). These frameworks propose that sapwood area at a given height is proportional to the branch sapwood and leaf area it supports, implying that conductive area at tap height reflects both the number and size of branches connected to the trunk. In turn, greater sapwood area should support higher sap flow and sap volume yields. Although these allometric rules have limitations (Lehnebach et al., 2018), the pipe model has been shown to perform well in a maple species (*Acer mono*), where it held across most branch orders, except for current-year branches (Sone et al., 2005). In sugar maple, our results are consistent with this framework: for a given DBH, trees with more branches along the trunk (longer crown length) and greater branch biomass relative to the stem tended to produce higher sap volumes.

### MAPLE SYRUP YIELDS

Crown surface area was the strongest predictor of maple syrup yield after controlling for DBH, which is encouraging from a practical standpoint, as it can be easily estimated in the field using simple measurements (e.g., longest crown diameter and a perpendicular diameter)(Schomaker et al., 2007). This makes it a feasible metric for inclusion in routine sugarbush surveys, alongside crown length that was highly influential on sap volume. In contrast, other variables examined in this study derived from LIDAR data, such as wood volume, wood surface area, or crown volume, are far more difficult to obtain without specialized equipment.

However, accurate estimation of both yield components (sap volume and sugar content) is not always necessary. Under high-vacuum systems, where sap yield from the same tree and tap can increase severalfold relative to gravity methods (Wilmot et al., 2007), sap sugar content appears to play a stronger role in syrup production, as shown in our results (36% sugar content and 54% sap volume). In contrast, our previous and ongoing work under bucket collection shows that syrup yield is driven primarily by sap volume, explaining over 90% of the variation (Bouchard et al. 2025; Bouchard et al., in prep), with little contribution from sugar content. Thus, harvesting systems may shift which variables are most important to predict inter- individual yields: high-vacuum systems require accurate estimation of both sap volume and sugar content, making crown area a useful proxy, whereas gravity systems should emphasize sap volume.

### CONCLUSIVE REMARKS AND MANAGEMENT IMPLICATIONS

The architectural traits that maximize maple syrup production contrast sharply with those typically desired in commercial forestry. In forestry, trees with long, branch-free stems are preferred, as they provide high- quality logs with minimal knots. In maple sugaring, however, our results showed that a greater number and size of branches along the trunk is beneficial. This fundamental difference underscores the importance to adapt forest management practices to the commercial context. Silvicultural treatments that promote crown development, such as thinning, appear promising in maple sugaring.

A previous study examining the effects of thinning on maple syrup yield and its components did not report any long-term benefits on yield 15 years after treatment (Pothier, 1995). This may be partly explained by the use of plot-level averages rather than individual tree yields, potentially masking tree-level variability, as well as by the relatively low vacuum level applied (≈0.07 MPa; ∼20 in Hg), which is below that of modern systems. In addition, the study focused on radial growth responses without assessing crown development, limiting evaluation of whether thinning achieved the expected effects on crown expansion. Therefore, future work should be needed to further investigate thinning approaches.

Overall, research on forest management in the context of maple syrup production remains scarce. By providing quantitative evidence on the effects of growth and tree and crown size and structure on yields in mature sugar bushes, our study provides a strong foundation for future research in this area.

## NUMBER OF WORDS

6336

## DATA AVAILABILITY STATEMENT

The data will be publicly available on Figshare upon publication.

## CONFLICT OF INTEREST

The authors declare no conflict of interest.

## ACKNOWLEDGEMENTS

This work was supported in part by a NSERC Doctoral Scholarship awarded to Élise Bouchard and a MITACS grant awarded to Gauthier Lapa. We thank Stéphane Corriveau for his technical assistance throughout the project.

## Notes

### Competing Interest Statement

The authors have declared no competing interest.

