## Supplementary material for "A crown story: How tree architecture drives maple syrup yields"

### APPENDIX S1 – Barrel Sanitization Protocol

**Pre-rinsing**

Rinse the barrels with hot water (40–45 °C) before cleaning.

- Use 20 L of hot water for approximately 1 minute.
- Drain the rinse water.

**Preparation of the cleaning solution**

- In a 20 L bucket, add hot water (40–45 °C) up to the upper mark (approximately 18 L).
- Add and mix 100 mL of Germac (quaternary amine) into the water.

**Barrel cleaning**

- Transfer the cleaning solution into the barrel.
- Close the barrel with the cap, allowing a few drops to escape to release pressure caused by the hot water.
- Using the barrel washer, rotate the barrel for 2 minutes.
- Drain the barrel contents.

**Final rinsing**

- Using a hose and spray nozzle, rinse the barrel with hot water (40–45 °C) until no visible cleaning residue or foam remains.
- Add 20 L of clean hot water to the barrel.
- Close the barrel with the cap, allowing a few drops to escape to release pressure caused by the hot water.
- Using the barrel washer, rotate the barrel for 1 minute.
- Drain the barrel contents.

**Barrel drying**

- Using wooden blocks, elevate the barrels in a tilted vertical position (cap facing upward) so that residual liquid collects on one side of the barrel bottom.
- Using a vacuum pump, remove the residual liquid through the 2-inch cap opening.
- Inject compressed air through the ¾-inch or 2-inch cap opening to facilitate drying.
- Using a flashlight, inspect the barrel for any remaining liquid. If necessary, repeat the drying steps.

**Barrel storage**

- Once the barrel is clean and dry, securely tighten the caps before storage.

### APPENDIX S2 - Model selection

SUGAR CONTENT

Model selection was performed first on a subset of 34 trees to include growth rate in the model selection, since growth data were missing for 4 trees. If the best model on this subset did not include growth rate, selection was repeated on the full dataset of 38 trees, excluding models with growth rate.

**Table S2.1** Sugar content models, associated hypotheses, and included variables. Wood volume of the main stem was included in all models to account for overall tree size, rather than DBH, due to its closer relationship with sap sugar content, but is omitted here for clarity.

| **Model** | **Hypothesis** | **Variables** |
|---|---|---|
| M1 | Crown | Crown area |
| M2 | Crown | Crown volume |
| M3 | Crown | Wood surface branches |
| M4 | Shape | Crown area + Crown asymmetry |
| M5 | Shape | Crown area + Fork rate |
| M6 | Distance | Crown area + Crown height |
| M7 | Stress | Crown area + Growth 25mm |

Model selection was conducted in sequential steps to identify the combination of variables that best explained sugar content.

1. **Selection of the crown size variable:** Three candidate models representing the Crown hypothesis were compared: crown area (M1), crown volume (M2), and branch wood surface (M3). Model M1, based on crown area, was selected as the best-performing model.
2. **Adding potential architectural modifiers:** Starting from M1, additional architectural variables were added to test the Shape hypothesis. Two models were considered: crown area with crown asymmetry (M4) and crown area with fork rate (M5). The two best models (M1 and M4) were equivalent ($\Delta$AICc < 3), therefore the most parsimonious model was selected (M1).
3. **Incorporating factors modulating the effect of crown size:** Finally, variables representing distance to the crown (M6) and recent growth as a stress indicator (M7) were added to M1. Model M7, which included crown area and recent growth, was selected as the best overall model, with a $\Delta$AICc of 11.04 relative to the second-best model (Table S2.2).

**Table S2.2** Model selection results for sugar content based on a subset of 34 trees for which growth rate data were available. Models were compared using Akaike Information Criterion (AICc). For each candidate model, the number of estimated parameters (K), log-likelihood (LL), corrected AICc, ΔAICc, Akaike weights (AICcWt), and cumulative Akaike weights (Cum.Wt) are reported.

| Model | K | AICc | Delta_AICc | AICcWt | Cum.Wt | LL |
|---|---|---|---|---|---|---|
| M7 | 5 | 40.87 | 0 | 0.99 | 0.99 | -14.36 |
| M4 | 5 | 51.91 | 11.04 | 0 | 1 | -19.88 |
| M1 | 4 | 54.48 | 13.61 | 0 | 1 | -22.55 |
| M6 | 5 | 55.92 | 15.05 | 0 | 1 | -21.89 |
| M5 | 5 | 56.64 | 15.77 | 0 | 1 | -22.25 |
| M3 | 4 | 60.12 | 19.25 | 0 | 1 | -25.37 |
| M2 | 4 | 60.39 | 19.52 | 0 | 1 | -25.5 |

SAP VOLUME

Model selection was performed first on a subset of 34 trees to include growth rate in the model selection, since growth data were missing for 4 trees. If the best model on this subset did not include growth rate, selection was repeated on the full dataset of 38 trees, excluding models with growth rate.

**Table S2.3** Sap volume models, associated hypotheses, and included variables. DBH was included in all models to account for overall tree size but is omitted here for clarity, except for Model 2, which included DBH as the sole predictor and is explicitly indicated in the table.

| **Model** | **Hypothesis** | **Variables** |
|---|---|---|
| M1 | Pressure | Wood surface area |
| M2 | Wood volume | DBH |
| M3 | Wood volume | Wood volume |
| M4 | Wood volume | Wood volume main axis + Wood volume branches |
| M5 | Combined | Wood volume main axis + Wood surface area |
| M6 | Branch | Crown volume |
| M7 | Branch | Crown area |
| M8 | Branch | Crown length |
| M9 | Branch | Crown length + Wood volume main axis + Wood volume branches |
| M10 | Growth | Growth 75 mm |
| M11 | Growth | Growth 75 mm + Crown volume |

| | | |
|---|---|---|
| M12 | Growth | Growth 75 mm + Wood volume main axis + Wood volume branches |

**Table S2.4** Model selection results for sap volume based on the full dataset (38 trees). Models were compared using AICc. For each candidate model, the number of estimated parameters (K), log-likelihood (LL), corrected Akaike Information Criterion (AICc), ΔAICc, and Akaike weights ($w_i$) are reported.

| Model | K | AICc | Delta_AICc | AICcWt C | Cum.Wt | LL |
|---|---|---|---|---|---|---|
| M9 | 6 | 334.7 | 0 | 0.53 | 0.53 | -160 |
| M8 | 4 | 336 | 1.29 | 0.28 | 0.81 | -163.4 |
| M6 | 4 | 338.3 | 3.58 | 0.09 | 0.89 | -164.5 |
| M4 | 5 | 339.4 | 4.75 | 0.05 | 0.94 | -163.8 |
| M5 | 5 | 340.3 | 5.61 | 0.03 | 0.98 | -164.2 |
| M1 | 4 | 341.6 | 6.91 | 0.02 | 0.99 | -166.2 |
| M7 | 4 | 343.7 | 9.05 | 0.01 | 1 | -167.3 |
| M2 | 3 | 346.2 | 11.49 | 0 | 1 | -169.7 |
| M3 | 4 | 347.8 | 13.08 | 0 | 1 | -169.3 |

Models 9 and 8 (branch hypothesis) were both well supported as best models (ΔAICc = 1.28). Model 8 is more parsimonious, including only two predictors, whereas Model 9 incorporates four. All predictors were statistically significant in both models.

MAPLE SYRUP

Model selection was performed first on a subset of 34 trees to include growth rate in the model selection, since growth data were missing for 4 trees. If the best model on this subset did not include growth rate, selection was repeated on the full dataset of 38 trees, excluding models with growth rate.

**Table S2.5** Maple syrup models, associated hypotheses, and included variables.

| **Model** | **Hypothesis** | **Variables** |
|---|---|---|
| M1 | Single component | Crown area + Growth 25mm + DBH |
| M2 | Single component | Crown length + DBH |
| M3 | Combined | Crown length + Crown area + DBH |
| M4 | Reduced | Crown area |
| M5 | Reduced | Crown area + DBH |
| M6 | Control | DBH |

**Table S2.6** Model selection results for maple syrup based on the full dataset (38 trees). Models were compared using AICc. For each candidate model, the number of estimated parameters (K), log-likelihood (LL), corrected Akaike Information Criterion (AICc), ΔAICc, and Akaike weights ($w_i$) are reported.

| Model | K | AICc | Delta_AICc | AICcWt | Cum.Wt | LL |
|---|---|---|---|---|---|---|
| M5 | 3 | 88 | 0 | 0.4 | 0.4 | -41 |
| M4 | 4 | 89 | 0.26 | 0.35 | 0.76 | -40 |
| M3 | 5 | 89 | 1.03 | 0.24 | 1 | -39 |
| M2 | 4 | 99 | 10.3 | 0 | 1 | -45 |
| M6 | 3 | 102 | 14.01 | 0 | 1 | -48 |

Models 4, 5, and 3, corresponding to the Combined and Reduced hypotheses, were well supported (ΔAICc ≤ 2.66). Model 5 was the most parsimonious (K = 3), but exhibited heteroscedasticity in residuals. Consequently, we selected model 4 as the best candidate model.

### APPENDIX S3 – Pairwise relationships

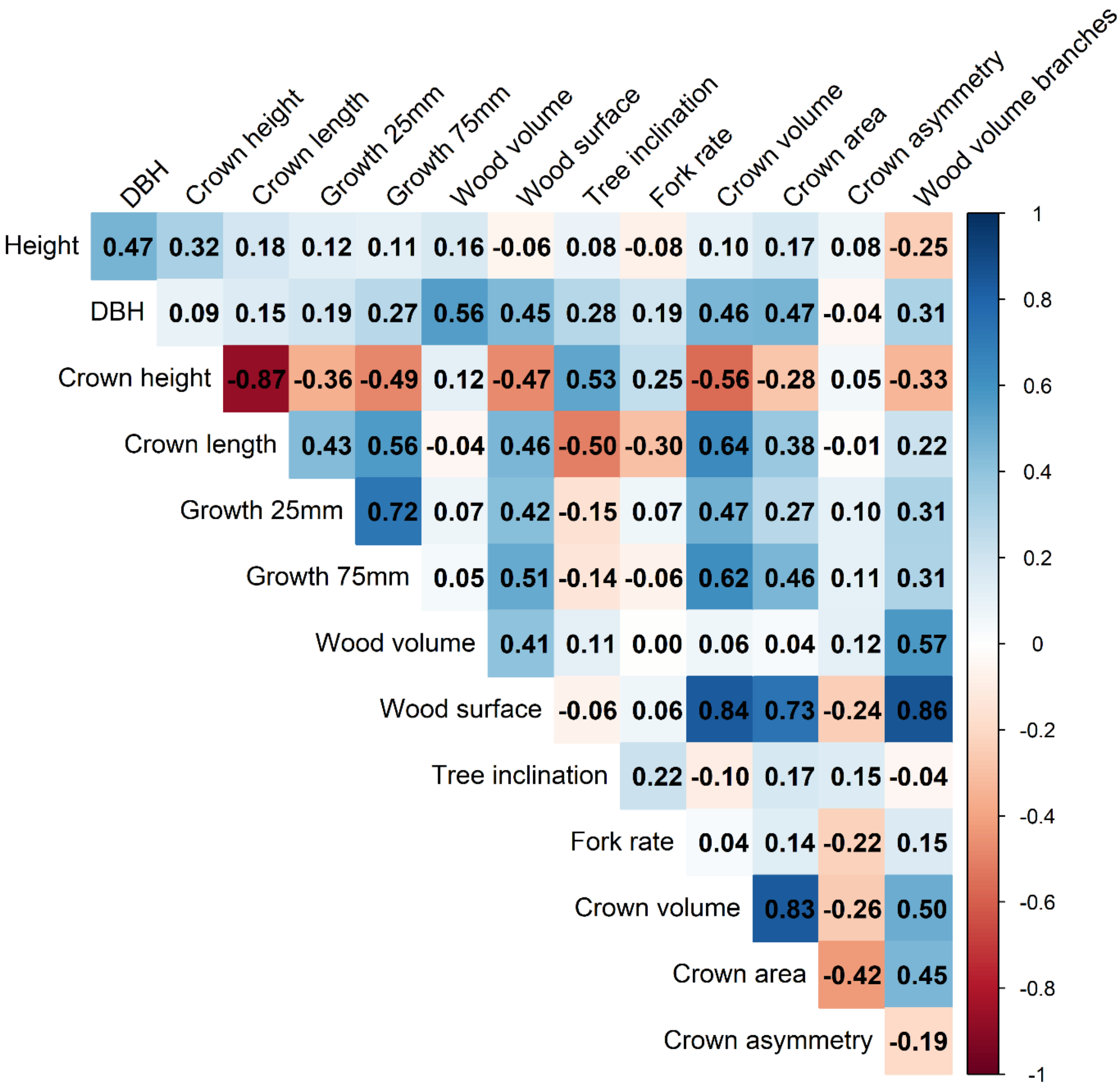


**Fig. S3.1** Pearson correlations (r) among tree architectural and growth variables.

**Table S3.1** Bivariate linear relationships between sap sugar content and morphological, growth, or structural variables. Columns show adjusted $R^2$ (adj$R^2$), p-value, model coefficient (Estimate), and standard error (Std. error). Grey cells indicate non-significant relationships (p-value > 0.05).

| | SAP SUGAR CONTENT | | | |
|---|---|---|---|---|
| Variable Name | adj$R^2$ | *p-value* | Estimate | Std. Error |
| Crown area | 0.17 | 0.01 | 0.02 | 0.01 |
| Crown asymmetry | 0.14 | 0.01 | -3.48 | 1.30 |

| | | | | |
|---|---|---|---|---|
| Growth rate 25 mm | 0.11 | 0.03 | -0.42 | 0.19 |
| Crown diameter | 0.09 | 0.03 | 0.16 | 0.07 |
| Wood surface branches | 0.09 | 0.03 | 0.01 | 0.00 |
| Wood surface | 0.09 | 0.04 | 0.01 | 0.00 |
| Wood volume | 0.08 | 0.04 | 0.29 | 0.14 |
| Wood volume main axis | 0.07 | 0.06 | 0.30 | 0.15 |
| Wood volume branches | 0.06 | 0.07 | 2.15 | 1.16 |
| Crown volume | 0.05 | 0.10 | 0.00 | 0.00 |
| DBH | 0.02 | 0.18 | 2.44 | 1.77 |
| Wood surface main axis | -0.01 | 0.39 | 0.02 | 0.02 |
| Growth rate 75 mm | -0.02 | 0.49 | -0.10 | 0.15 |
| Tree inclination | -0.02 | 0.60 | 0.78 | 1.48 |
| Fork rate | -0.03 | 0.77 | -0.01 | 0.03 |
| Height | -0.03 | 0.78 | 0.02 | 0.07 |
| Crown height | -0.03 | 0.78 | 0.01 | 0.03 |
| Crown length | -0.03 | 0.89 | 0.00 | 0.03 |

**Table S3.2** Bivariate linear relationships between sap volume and morphological, growth, or structural variables. Columns show adjusted $R^2$ (adj$R^2$), p-value, model coefficient (Estimate), and standard error (Std. error). Grey cells indicate non-significant relationships (p-value > 0.05).

| | **SAP VOLUME** | | | |
|---|---|---|---|---|
| Variable Name | adj$R^2$ | *p-value* | Estimate | Std. error |
| Crown volume | 0.32 | 0.00 | 0.11 | 0.03 |
| Crown length | 0.27 | 0.00 | 4.75 | 1.24 |
| Wood surface | 0.24 | 0.00 | 0.36 | 0.10 |
| Wood surface branches | 0.21 | 0.00 | 0.37 | 0.11 |
| Crown area | 0.21 | 0.00 | 0.86 | 0.26 |
| Growth rate 75 mm | 0.19 | 0.01 | 15.83 | 5.31 |
| Wood surface main axis | 0.19 | 0.00 | 2.60 | 0.85 |
| Crown height | 0.17 | 0.01 | -3.68 | 1.26 |
| Growth rate 25 mm | 0.16 | 0.01 | 19.95 | 7.29 |
| DBH | 0.15 | 0.01 | 190.24 | 69.82 |
| Wood volume branches | 0.13 | 0.01 | 121.83 | 47.02 |
| Crown diameter | 0.09 | 0.04 | 6.69 | 3.12 |
| Height | 0.00 | 0.37 | 2.55 | 2.81 |
| Tree inclination | -0.01 | 0.47 | -45.41 | 62.37 |
| Wood volume | -0.02 | 0.55 | 3.80 | 6.28 |
| Wood volume main axis | -0.02 | 0.75 | 2.20 | 6.75 |
| Fork rate | -0.03 | 0.83 | 0.23 | 1.11 |
| Crown asymmetry | -0.03 | 0.97 | 2.21 | 59.94 |

**Table S3.3** Bivariate linear relationships between maple syrup yield and morphological, growth, or structural variables. Columns show adjusted $R^2$ ($adjR^2$), p-value, model coefficient (Estimate), and standard error (Std. error). Grey cells indicate non-significant relationships (p-value > 0.05).

| | **MAPLE SYRUP YIELD** | | | |
|---|---|---|---|---|
| Variable Name | $adjR^2$ | *p-value* | Estimate | Std. error |
| Crown area | 0.45 | 0.00 | 0.05 | 0.01 |
| Crown volume | 0.41 | 0.00 | 0.01 | 0.00 |
| Wood surface | 0.39 | 0.00 | 0.02 | 0.00 |
| Wood surface branches | 0.36 | 0.00 | 0.02 | 0.00 |
| Wood volume branches | 0.24 | 0.00 | 6.61 | 1.85 |
| Crown diameter | 0.23 | 0.00 | 0.42 | 0.12 |
| DBH | 0.21 | 0.00 | 9.28 | 2.83 |
| Wood surface main axis | 0.19 | 0.00 | 0.11 | 0.04 |
| Crown length | 0.14 | 0.01 | 0.15 | 0.06 |
| Growth rate 75 mm | 0.07 | 0.07 | 0.46 | 0.24 |
| Crown height | 0.07 | 0.06 | -0.11 | 0.06 |
| Wood volume | 0.06 | 0.08 | 0.45 | 0.25 |
| Crown asymmetry | 0.05 | 0.10 | -4.11 | 2.42 |
| Wood volume main axis | 0.03 | 0.15 | 0.40 | 0.28 |
| Height | -0.01 | 0.38 | 0.11 | 0.12 |
| Growth rate 25 mm | -0.02 | 0.52 | 0.22 | 0.34 |
| Tree inclination | -0.03 | 0.79 | -0.71 | 2.64 |
| Fork rate | -0.03 | 0.96 | 0.00 | 0.05 |
